# Application of a novel haplotype-based scan for local adaptation to study high-altitude adaptation in rhesus macaques

**DOI:** 10.1101/2020.05.19.104380

**Authors:** Zachary A. Szpiech, Taylor E. Novak, Nick P. Bailey, Laurie S. Stevison

**Affiliations:** Department of Biology Pennsylvania State University University Park, PA 16801 USA; Institute for Computational and Data Sciences Pennsylvania State University University Park, PA 16801 USA; Department of Biological Sciences Auburn University Auburn, AL 36842, USA

**Keywords:** Adaptation, genome-scan, high-altitude, EGLN1, selection, macaque

## Abstract

When natural populations split and migrate to different environments, they may experience different selection pressures that can lead to local adaptation. To capture the genomic patterns of a local selective sweep, we develop XP-nSL, a genomic scan for local adaptation that compares haplotype patterns between two populations. We show that XP-nSL has power to detect ongoing and recently completed hard and soft sweeps, and we then apply this statistic to search for evidence of adaptation to high altitude in rhesus macaques. We analyze the whole genomes of 23 wild rhesus macaques captured at high altitude (mean altitude > 4000m above sea level) to 22 wild rhesus macaques captured at low altitude (mean altitude < 500m above sea level) and find evidence of local adaptation in the high-altitude population at or near 303 known genes and several unannotated regions. We find the strongest signal for adaptation at EGLN1, a classic target for convergent evolution in several species living in low oxygen environments. Furthermore, many of the 303 genes are involved in processes related to hypoxia, regulation of ROS, DNA damage repair, synaptic signaling, and metabolism. These results suggest that, beyond adapting via a beneficial mutation in one single gene, adaptation to high altitude in rhesus macaques is polygenic and spread across numerous important biological systems.

**Impact Summary:** When positive selection is ongoing or a beneficial mutation has recently fixed in a population, genetic diversity is reduced in the vicinity of the adaptive allele, and we expect to observe long homozygous haplotypes at high frequency. Here we develop a statistic that summarizes these expected patterns and compares between two populations in order to search for evidence of adaptation that may have occurred in one but not the other. We implement this statistic in a popular and easy-to-use software package, and then apply it to study adaptation to high altitude in rhesus macaques.

Extreme environments pose a challenge to life on multiple fronts. Very high-altitude environments are one such example, with low atmospheric oxygen, increased ultraviolet light exposure, harsh temperatures, and reduced nutrition availability. In spite of these challenges, many plants and animals, including humans, have genetically adapted to cope with these hardships. Here we study two populations of rhesus macaques, one living at high altitude and one living close to sea level. We apply our novel statistic to compare their haplotype patterns between them to search for evidence of genetic changes that are indicative of adaptation to their environment.

We find evidence for adaptation at a critical gene that helps control physiological response to low-oxygen, one that has been the target of repeated convergent evolution across many species. We also find evidence for positive selection across a range of traits, including metabolic and neurological. This work helps to explain the evolutionary history of the rhesus macaque and furthers our understanding about the ways organisms genetically adapt to high-altitude environments.

## Introduction

Selective sweeps produce regions of reduced genetic diversity in the vicinity of an adaptive mutation. These patterns manifest as long extended regions of homozygous haplotypes segregating at high frequency (Przeworski 2002; Sabeti *et al*. 2002; Kim and Nielsen 2004; Garud *et al*. 2015). In the event of a *de novo* mutation that is adaptive in a population, we expect the haplotype it resides on to rapidly rise in frequency in the population (called a ‘hard’ sweep). On the other hand, if an ancestrally segregating neutral or mildly deleterious allele turned out to be adaptive in a new environment, it would likely reside on two or more haplotypes, which would rapidly rise in frequency in the population (called a ‘soft’ sweep) (Hermisson and Pennings 2005; Pennings and Hermisson 2006). As both of these processes happen on a time scale faster than mutation or recombination can act to break up the sweeping haplotypes, we expect to observe long and low diversity haplotypes at high frequency in the vicinity of an adaptive mutation. However, if this mutation either does not exist or is not adaptive in a sister population, we would not expect a sweep to occur and thus we would not expect to observe similar haplotype patterns.

To capture these haplotype patterns and contrast them between a pair of populations, we develop XP-nSL, a haplotype-based statistic with good power to detect partial, fixed, and recently completed hard and soft sweeps by comparing a pair of populations. XP-nSL is an extension of nSL (Ferrer-Admetlla *et al*. 2014) and does not require a genetic recombination map for computation. The lack of dependence on a recombination map is important, as other statistics for identifying positive selection are biased towards low-recombination regions (O’reilly *et al*. 2008), but the approach taken by nSL has been shown to be more robust (Ferrer-Admetlla *et al*. 2014). Both nSL and XP-nSL summarize haplotype diversity by computing the mean number of sites in a region that are identical-by-state across all pairs of haplotypes. Whereas nSL contrasts between haplotype sets carrying an ancestral or a derived allele in a single population, XP-nSL contrasts between haplotype sets in two different populations, allowing it to detect local adaptation.

An extreme example of adaptation to a local environment is the transition to high-altitude living. Organisms living at high altitude are confronted with many challenges, including a low-oxygen atmosphere and increased ultraviolet light exposure, and these harsh environments inevitably exert strong selection pressure. Indeed, adaptation to high-altitude living has been studied extensively across many organisms from plants, including monocots (Gonzalo-Turpin and Hazard 2009; Ahmad *et al*. 2016) and dicots (Kim and Donohue 2013; Liu *et al*. 2014; Munne-Bosch *et al*. 2016; Guo *et al*. 2018), to numerous animals including amphibians (Yang *et al*. 2016), canids (Li *et al*. 2014; Wang *et al*. 2014; Wang *et al*. 2020), humans (Bigham *et al*. 2009; Bigham *et al*. 2010; Xu *et al*. 2010; Yi *et al*. 2010; Peng *et al*. 2011; Huerta-Sanchez *et al*. 2013; Huerta-Sanchez *et al*. 2014; Jeong *et al*. 2014), yaks (Qiu *et al*. 2012), birds (Cai *et al*. 2013; Qu *et al*. 2013; Wang *et al*. 2015; Graham and Mccracken 2019), boars (Li *et al*. 2013), mice (Storz *et al*. 2007; Cheviron *et al*. 2012; Schweizer *et al*. 2019; Storz *et al*. 2019; Velotta *et al*. 2020), moles (Campbell *et al*. 2010), antelope (Ge *et al*. 2013), and horses (Hendrickson 2013). Liu *et al*. (2018) recently sequenced and published the whole genomes of 79 wild-born Chinese rhesus macaques collected from multiple sites in China. Among these animals, 23 were sampled from far western Sichuan province in a region with mean altitude > 4000 m above sea level (Liu *et al*. 2018), providing an opportunity to study the genetics of local adaption to high altitude in rhesus macaques.

Rhesus macaques are the second most widely distributed primate, with a range extending from Afghanistan to Vietnam and from a latitude of 15 to 38 degrees north (Fooden 2000). Early ancestors of the macaque migrated out of Africa to the Eurasian continent approximately 7 mya—the earliest catarrhine fossils on the continent are macaque-like (Stewart and Disotell 1998). Modern rhesus macaques trace a recent origin to Southeast Asia, with a major migratory split occurring approximately 162 kya separating the ancestors of modern Indian and Chinese rhesus macaques (Hernandez *et al*. 2007). Macaques have proven to be quite evolutionarily successful, demonstrating ecological flexibility and adaptability via developmental plasticity and behavioral changes (Richard *et al*. 1989; Madrid *et al*. 2018). Other studies have looked at how the rhesus macaque radiation has led to population-level adaptation to climate and food availability (Liu *et al*. 2018).

Here we test and evaluate our XP-nSL statistic and apply it to study the genomic consequences of high-altitude living in the rhesus macaque. We use it to compare the haplotype patterns of the 23 animals from the high-altitude population with another 22 that Liu *et al*. (2018) sampled in lower-lying regions in eastern China with a mean altitude < 500 m above sea level.

## Methods

### Data Preparation

Liu *et al*. (2018) generated whole genome sequencing data for 79 Chinese macaques and called all biallelic polymorphic sites according to GATK best practices using rheMac8, identifying 52,534,348 passing polymorphic autosomal sites. We then filter all loci with > 10% missing data leaving 35,639,395 biallelic sites. Next, the program SHAPEIT v4.1.2 (Delaneau *et al*. 2019) was used to phase haplotypes in the full data set with a genetic map that was available for rheMac8 (Bcm-Hgsc 2020). SHAPEIT performs imputation during phasing for any missing genotypes. Our analyses here focus on 45 of the 79 samples from Liu *et al*. (2018), representing 23 from high-altitude regions of China (*M. m. lasiotis*) and 22 from low-altitude regions of China (*M. m. littoralis*), based on capture location information. See Table S1 for individual IDs used.

Liu *et al*. (2018) also inferred joint demographic histories for their five populations, and we extract the demographic parameters for our two of interest. This demographic history is recapitulated in Fig. 1 with detailed parameters given in Table 1, which are then used for simulations to test XP-nSL.

**Figure 1.**
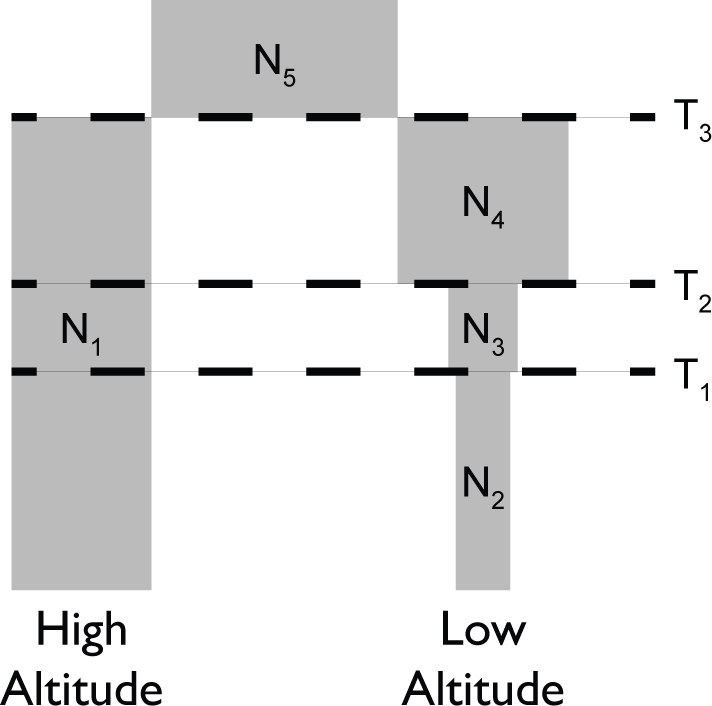
A representation of the demographic history for our high-and low-altitude populations as inferred by Liu *et al*. (2018).

**Table 1.**
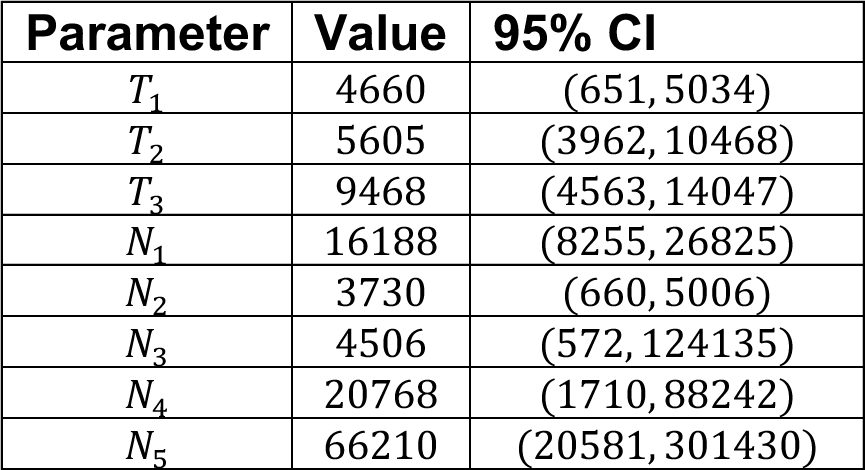
Demographic parameters used for simulations with 95% confidence intervals from (Liu *et al*. 2018). T values are given in number of generations before present. N values represent diploid effective population size.

| Parameter | Value | 95% CI |
| --- | --- | --- |
| $T_1$ | 4660 | (651, 5034) |
| $T_2$ | 5605 | (3962, 10468) |
| $T_3$ | 9468 | (4563, 14047) |
| $N_1$ | 16188 | (8255, 26825) |
| $N_2$ | 3730 | (660, 5006) |
| $N_3$ | 4506 | (572, 124135) |
| $N_4$ | 20768 | (1710, 88242) |
| $N_5$ | 66210 | (20581, 301430) |

### A Statistic for Detecting Local Adaptation

We developed a cross-population haplotype-based statistic, XP-nSL, to scan for regions of the genome implicated in local adaptation between two populations by extending nSL (Ferrer-Admetlla *et al*. 2014), each of which is defined below.

Consider the sets *A*(*k*) and *D*(*k*), representing the set of haplotypes at site *k* carrying the ancestral or derived allele, respectively, and let *n_A_*(*k*) = |*A*(*k*)| and *n_D_*(*k*) = |*D*(*k*)|. Next *L_ij_*(*k*) is defined as the number of consecutive sites at which haplotype *i* and *j* are identical-by-state (IBS) in the interval containing site *k*. Then nSL at site *k* is 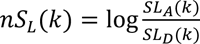, where 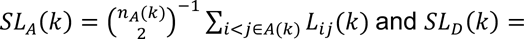 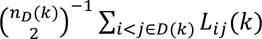. *SL*_*A*_(*k*) and *SL*_*D*_(*k*) represent the mean *L_ij_*(*k*) over all pairs of haplotypes carrying either the ancestral or derived allele at locus k, respectively. nSL scores are then normalized genome-wide in site-frequency bins either with respect to the empirical background or neutral simulations with a matching demographic history. The nSL computation is illustrated in Ferrer-Admetlla *et al*. (2014). nSL is implemented in `nsl` (Ferrer-Admetlla *et al*. 2014) and `selscan` v1.1.0+ (Szpiech and Hernandez 2014).

XP-nSL is defined similarly, except instead of comparing sets of haplotypes containing ancestral or derived alleles, it compares sets of haplotypes between two different populations. Let *P*_1_(*k*) and *P*_2_(*k*), represent the set of haplotypes at site *k* in population 1 and population 2, respectively, and 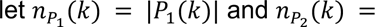 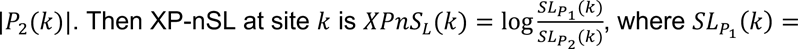 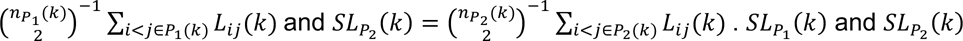 represent the mean *L*_ij_(*k*) over all pairs of haplotypes in population 1 or population 2 at locus k, respectively. XP-nSL scores are then normalized genome-wide either with respect to the empirical background or neutral simulations with a matching demographic history. The XP-nSL computation is illustrated in Fig. 2 with a toy example. We implement XP-nSL in `selscan` v1.3.0+ (Szpiech and Hernandez 2014) to facilitate wide adoption. It is worth noting that XP-nSL is analogous to XP-EHH (Sabeti *et al*. 2007) as nSL is analogous to iHS (Voight *et al*. 2006).

**Figure 2.**
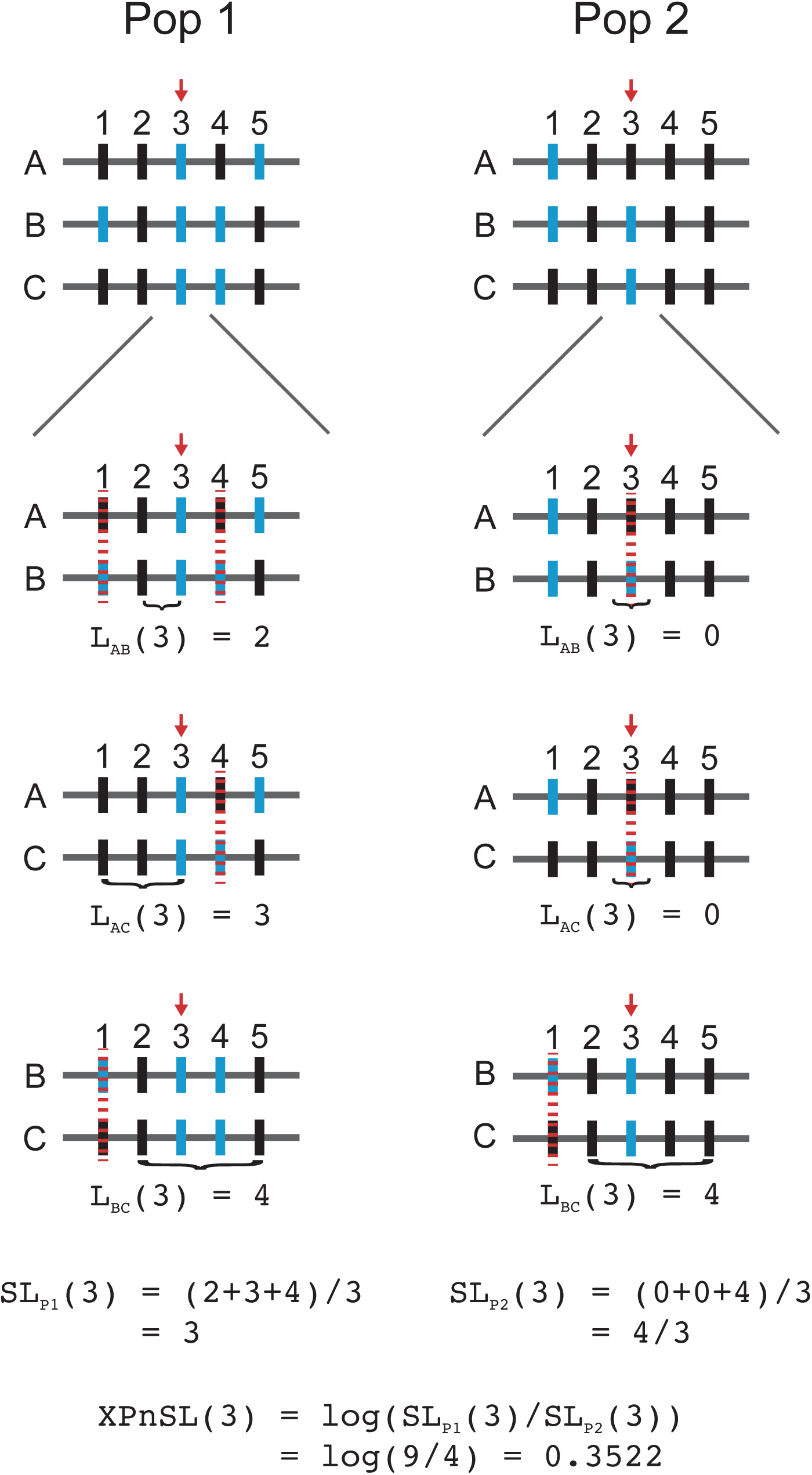
A toy example illustrating the computation of XP-nSL at a single site in two populations with three haplotypes (grey horizontal bars labeled A-C) and five sites (vertical bars labeled 1-5) where different alleles are colored blue or black. XP-nSL is calculated at site 3 (marked by red arrow). In each population, for each pair of haplotypes, the number of identical-by-state (IBS) sites are counted extending out from and including the test site (red arrow) until reaching a non-IBS site (marked by red dotted line). Within each population, the mean number of IBS sites is calculated across all pairs of haplotypes, and then the log-ratio of the mean from each population is computed to get XP-nSL at site 3.

The goal of these statistics is to capture a signal of extended regions of low diversity on sweeping haplotypes (emblematic of an ongoing or recently completed selective sweep) within a population (nSL) or on sweeping haplotypes in one population versus another (XP-nSL). When XP-nSL scores are positive this suggests evidence for a hard or soft sweep in population 1, and when XP-nSL scores are negative this suggests evidence for a hard or soft sweep in population 2.

### Simulations

In order to test the ability of XP-nSL to detect ongoing and recently completed hard and soft sweeps, coalescent simulations were performed conditional on an allele frequency trajectory with the program `discoal` (Kern and Schrider 2016). `discoal` simulates an allele frequency trajectory for a single non-neutral allele backwards in time and then simulates a neutral coalescent process conditional on this trajectory. This takes advantage of the speed and efficiency of the coalescent while still being able simulate genetic diversity patterns in the vicinity of a non-neutral locus.

All simulations were run with a two-population divergence demographic history (Fig. 1 and Table 1), as inferred by Liu *et al*. (2018), and given by the following `discoal` command line arguments `-p 2 46 44 -en 0 1 0.230410476572876 - en 0.071964666275442 1 0.278345739259351 -en 0.086558359329153 1 1.282885999320505 -ed 0.146214905642895 1 0 -en 0.146214905642895 0 4.089940389782871`. Here 46 haplotypes were sampled from population 0, which has the demographic history of the high-altitude population, and 44 haplotypes were sampled from population 1, which has the demographic history of the low-altitude population. A mutation rate of *μ* = 2.5 × 10^-8^ (Fan *et al*. 2018) was used, along with a recombination rate of *r* = 5.126 × 10^-9^, which was computed as the genome-wide mean rate from the rheMac8 recombination map (Bcm-Hgsc 2020). A 500 kb region was simulated, thus giving a scaled mutation rate and scaled recombination parameters for `discoal` as `-t 809.425 -r 165.967`. 5,349 replicates of neutral sequence were simulated under this model, representing approximately the entire macaque genome minus 500 kb. Thus, the total simulated length of all neutral regions plus one selected region is approximately equal to the macaque genome length.

For non-neutral simulations, sweep scenarios were simulated with a positive additive selection coefficient *s* ∈ {0.01,0.02,0.05}, which is provided to `discoal` as a scaled selection coefficient 2*N*_1_*s* ∈ {323.77, 647.54, 1618.85} (`discoal` flag `-a`). Soft sweeps are simulated as a mutation that arose neutral and turned beneficial at a particular establishment frequency *e* ∈ {0.01,0.02,0.03,0.04,0.05,0.10,0.20} (`discoal` flag `-f`). Hard sweeps are simulated from a *de novo* mutation that was never neutral, i.e. *e* = 0. Finally, sweeps were conditioned on having either reached a certain frequency in the population at the time of sampling, *f* ∈ {0.7, 0.8, 0.9, 1.0} (`discoal` flag `-c`), or that the adaptive mutation reached fixation some number of generations prior to sampling, *g* ∈ {50, 100, 200}, which is provided to `discoal` in coalescent units *g*/2*N*_1_ ∈ {1.5443 × 10^-3^, 3.0886 × 10^-3^, 6.1774 × 10^-3^} (`discoal` flag `-ws`). For the sake of being conservative in our estimation of power (Voight *et al*. 2006; Sabeti *et al*. 2007), it was assumed the actual adaptive mutation remains unsampled (`discoal` flag `-h`) even though whole genome sequencing data are being analyzed. For each combination of parameter values, 500 replicates were simulated.

### Neutral Simulations of Mismatched Histories

We also generate neutral simulations for three mismatched demographic histories, in order to study how a mismatched demographic history may influence the power and false positive rates of XP-nSL if used as a normalization baseline. We name these mismatched histories “Rand”, “Under”, and “Over” and generate 5,349 replicates for each one.

For the “Rand” history, each parameter is (uniformly) randomly chosen from within the 95% CI as inferred by (Liu *et al*. 2018) (see Table 1). The parameters are resampled for each replicate of the “Rand” history, and the condition *T*_1_ < *T*_2_ < *T*_3_ is enforced. For the “Under” history, the present-day population size parameters *N*_1_ and *N*_2_ are the only ones modified. They are set to *N*_1_ = 8255 at the extreme low end of the 95% CI and *N*_2_ = 5006 at the extreme high end of the 95% CI. This represents a scenario where the difference in population sizes is underestimated. For the “Over” history, once again the present-day population size parameters N1 and N2 are the only ones modified. They are set to *N*_1_ = 26825 at the extreme high end of the 95% CI and *N*_2_ = 660 at the extreme low end of the 95% CI. This represents a scenario where the difference in population sizes is overestimated.

### Detecting Local Adaptation in Real Data

From the phased data set, animals captured at high altitude (n = 23) and animals captured at low altitude (n = 22) were subset (see Table S1). Using `selscan` v1.3.0 (Szpiech and Hernandez 2014) to compute raw XP-nSL scores across the genome (`selscan` flags –xpnsl –vcf high-altitude.vcf –vcf-ref low-altitude.vcf), scores were then normalized using the genome-wide empirical background with `selscan`’s `norm` v1.3.0 program (`norm` flag --xpnsl). The low-altitude population was used as the reference population, so positive XP-nSL scores correspond to long homozygous haplotypes and a possible sweep in the high-altitude population compared to the low-altitude population, and vice versa for negative XP-nSL scores. A Manhattan plot of genome-wide normalized XP-nSL scores > 2 is plotted in Fig. S7.

In order to identify regions implicated as potentially adaptive, we search for clusters of extreme scores along a chromosome. Using `selscan`’s companion program `norm v1.3.0`, the genome is divided into non-overlapping 100kb regions and both the maximum XP-nSL score and the fraction of XP-nSL scores > 2 are computed (`norm` flags –xpnsl –bp-win –winsize 100000). `norm` then creates 10 quantile bins (--qbins 10) for windows with more than 10 sites per window (--min-snps 10) and identifies the top 1% of windows with the highest fraction of extreme scores (Fig. S6 and Table S4). Each window is then annotated with the ensembl rheMac8 gene list, and a maximum XP-nSL score is assigned to a given gene based on the max-score in the 100 kb window with which it overlaps. If a gene overlaps more than one 100 kb window, it is assigned the top max-score from among the windows.

## Results

### Power Analysis of XP-nSL

First, we evaluate the performance of XP-nSL based on simulations. After computing XP-nSL for all sites in all simulations, scores were normalized by subtracting the mean and dividing by standard deviation of the neutral simulations, giving the neutral scores an approximately N(0,1) distribution (Fig. S1).

We consider the maximum score in a 100 kb interval as way to identify regions under positive selection similar to Voight *et al*. (2006). To get the null distribution of max-scores, the maximum score is computed in the central 100 kb of each neutral simulation. The distribution of max-scores in neutral simulations had a median of 2.093 with 95% of the mass between 0.804 and 3.492, which is represented in Fig. 3 as a solid horizontal black line (median) and two dashed horizontal black lines (95% interval). Next, the maximum score is computed in the central 100 kb of all non-neutral simulations, and the median and 95% intervals are plotted for each parameter combination. Fig. 3 shows good separation between the neutral distribution of max-scores and the distribution of max-scores for a range of non-neutral parameters, suggesting that our statistic can distinguish between neutral and non-neutral scenarios. Note that soft sweeps that start at 0.1 or 0.2 frequency see the least separation from the neutral distribution. To evaluate the power of the max-score statistic, the 99^th^ percentile of the max-score distribution is computed in the neutral distribution (*neutral*_99_ = 3.792) and power is calculated as the mass of each non-neutral max score distribution > *neutral*_99_. With this approach, even if the entire genome is neutrally evolving, at most 1% of the genome will be identified as putatively under selection thus fixing the false positive rate at 1% at most. Results are plotted in Fig. 4A, which shows good power to detect both incomplete and completed sweeps. For soft sweeps that start at a frequency ≤ 0.03, power is > 75% when the sweep is near or just past fixation; the ability to detect soft sweeps falls off for sweeps that start > 0.03 frequency.

**Figure 3.**
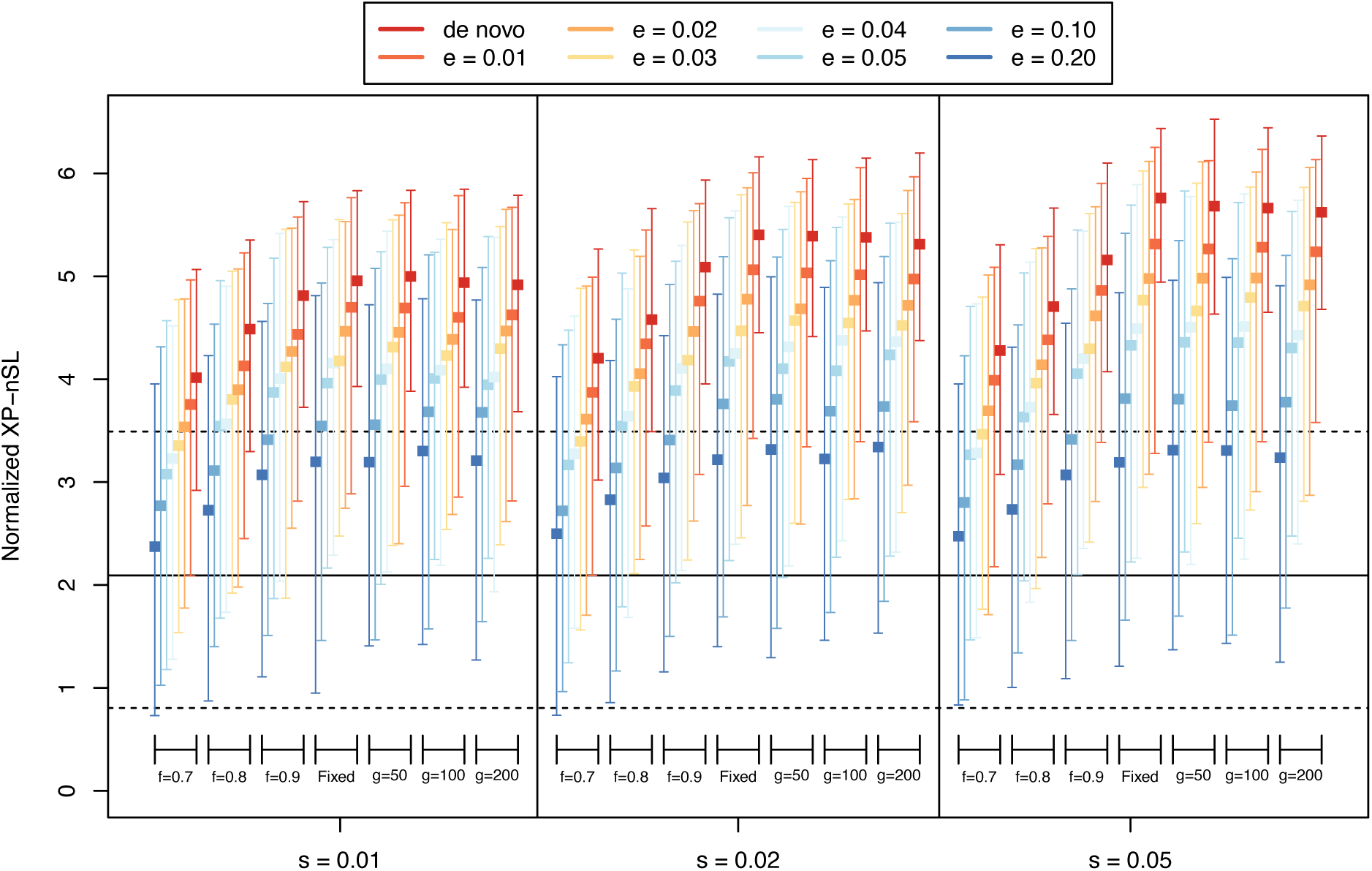
The distribution of maximum XP-nSL scores from simulations across various parameters, represented by medians and intervals containing 95% of the mass of the distribution. Neutral simulations are represented by the black solid horizonal line (median) and the black dashed horizonal line (95% interval). Non-neutral simulations represented by a colored box (median) and error bars (95% interval). The parameters are e (frequency at which selection begins, e > 0 indicates soft sweep), f (frequency of selected mutation at sampling), g (number of generations since fixation), and s (selection coefficient).

**Figure 4.**
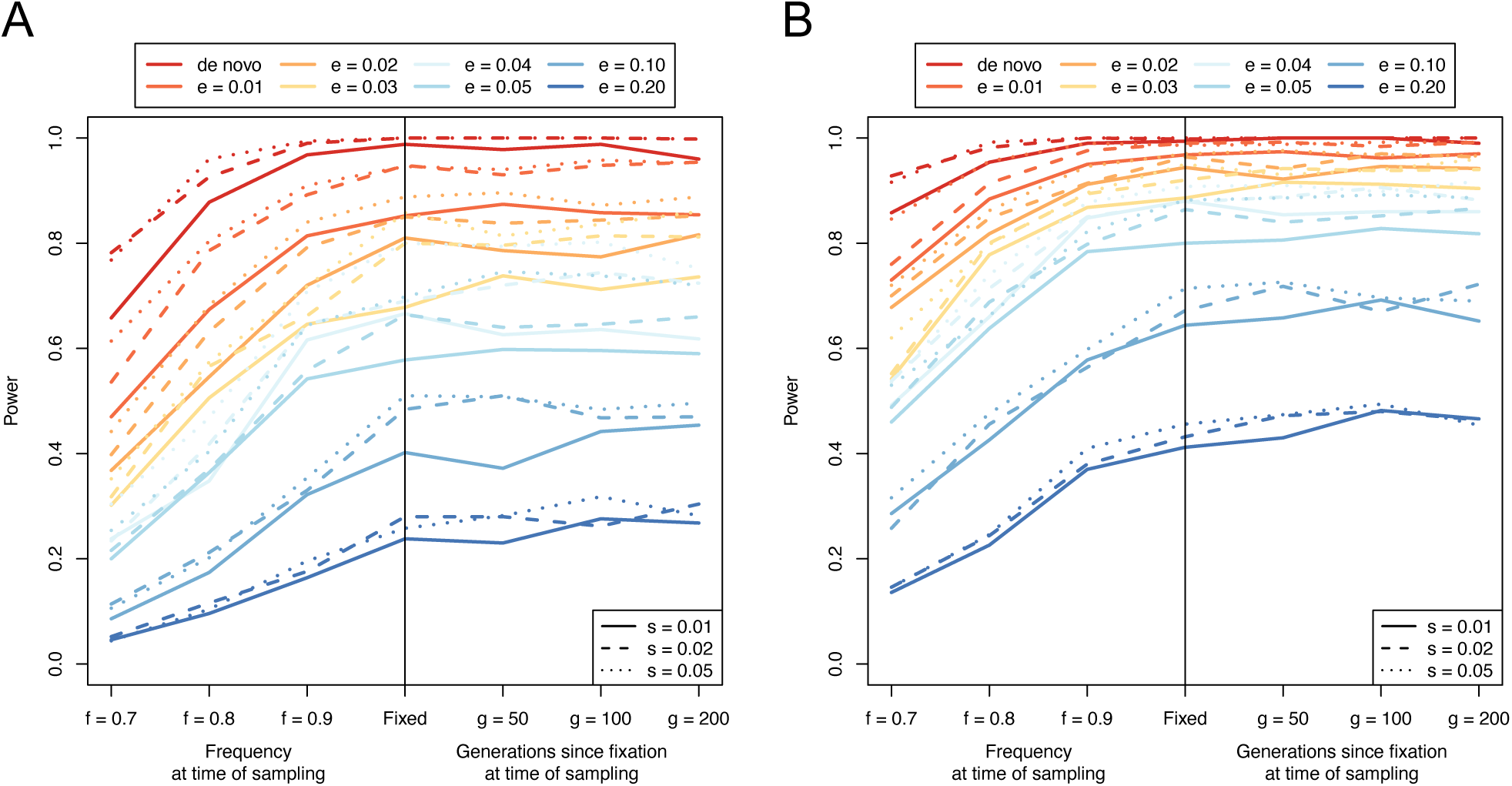
Power curves for (A) the max-score approach and (B) the window-based approach to identifying sweeps. The parameters are e (frequency at which selection begins, e > 0 indicates soft sweep), f (frequency of selected mutation at sampling), g (number of generations since fixation), and s (selection coefficient).

Next, we consider that due to linkage disequilibrium consecutive scores will be correlated, and we should therefore expect clusters of extreme scores in true non-neutral regions. We thus consider a window-based approach to identify selected regions similar to Voight *et al*. (2006), where the top 1% windows with a high number of extreme scores are identified. Taking the central 100 kb region of each simulation, the fraction of XP-nSL scores > 2 (representing approximately the highest 2% of all neutral scores) is computed. Since each 100 kb window has a variable number of sites within it, windows with fewer sites are more likely to have a higher fraction of extreme scores by chance. Thus, windows are binned by number of sites into 10 quantile bins, and the top 1% of windows with the highest fraction of extreme scores in each bin establishes the threshold beyond which a window is taken as putatively selected, as in Voight *et al*. (2006). With this approach, even if the entire genome is neutrally evolving, at most 1% of the genome will be identified as putatively under selection thus fixing the false positive rate at 1% at most. Power is computed for each non-neutral parameter set as the proportion of replicates for which the central 100 kb exceeds the 1% threshold as calculated from neutral simulations. The results are plotted in Fig. 4B, which shows improved power over the max-score approach across a wider range of parameters. Indeed, using the window-based method, power to detect soft sweeps improves substantially across the parameter space, with > 75% power to detect soft sweeps at or near fixation that started at frequency ≤ 0.05.

We next consider how XP-nSL power compares to nSL, XP-EHH, and F_ST_. We compare to nSL since XP-nSL is an extension of it, to XP-EHH as it is a similar haplotype-based two-population selection statistic, and to F_ST_ as it a popular two-population method used to infer local adaptation. For all simulations, nSL and XP-EHH are computed using `selscan` v1.3.0 (Szpiech and Hernandez 2014). Normalization, identification of top windows, and power calculation was done as described above for XP-nSL. F_ST_ is computed using `VCFtools` v0.1.16 (Danecek *et al*. 2011), which implements Weir and Cockerham’s formulation (Weir and Cockerham 1984), in 100kb windows for all simulations. The 99^th^ percentile F_ST_ value was determined from the central 100kb window among all neutral simulations. Power was then computed for each non-neutral parameter set as the proportion of replicates for which the F_ST_ value of the central 100kb window is greater than the neutral threshold as calculated from neutral simulations.

Fig. 5 shows the difference in power between XP-nSL and each statistic (raw power for nSL, XP-EHH, and F_ST_ shown in Fig. S4) over the simulated parameter space, where positive values indicate that XP-nSL has more power than the comparison statistic. XP-nSL is compared to nSL in Fig. 5A, which shows XP-nSL improves on nSL across nearly the entire parameter space and especially for sweeps that have fixed in the recent past. XP-nSL is compared to XP-EHH in Fig. 5B, which shows XP-nSL improving on XP-EHH for soft sweeps (*e* > 0). Finally, XP-nSL is compared to F_ST_ in Fig. 5C, which shows XP-nSL improving on F_ST_ mostly for incomplete sweeps, although F_ST_ performed better post-fixation for certain soft sweep scenarios (*e* ≥ 0.05).

**Figure 5.**
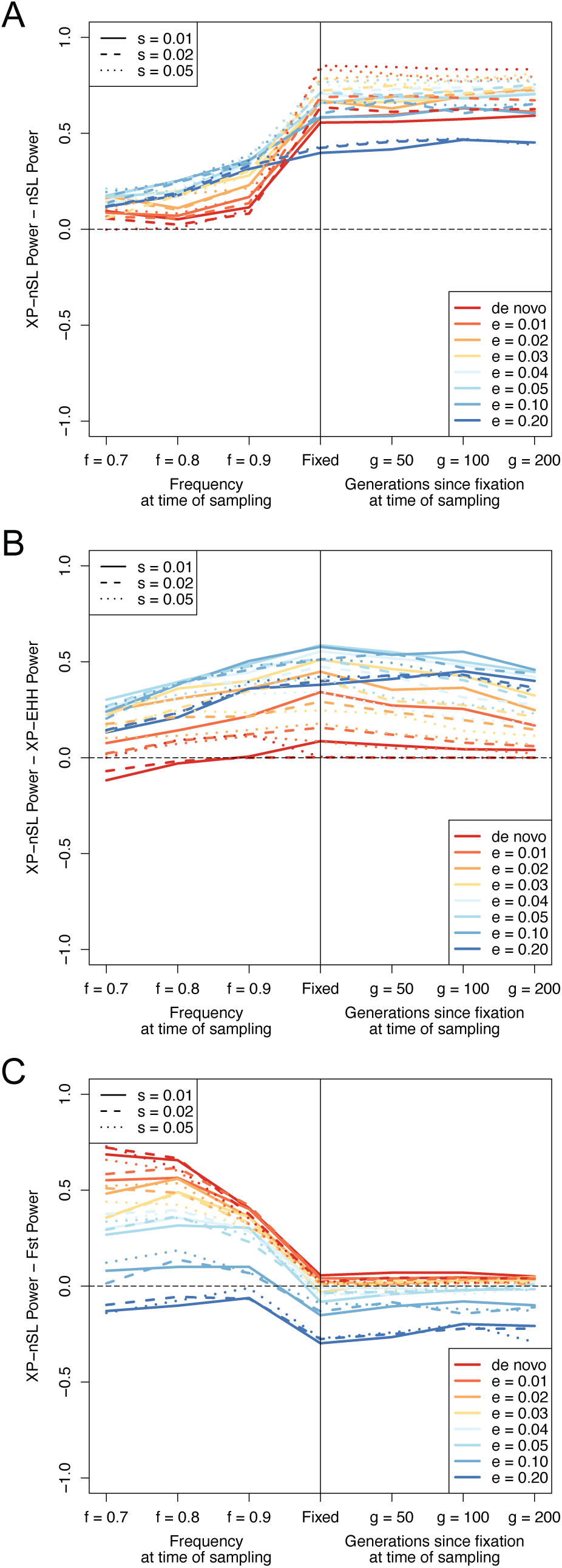
Difference in power between XP-nSL and (A) nSL, (B) XP-EHH, and (C) F_ST_. Values above 0 indicate XP-nSL has more power, and values below 0 indicate XP-nSL has less power. The horizontal black dotted line marks 0. The parameters are e (frequency at which selection begins, e > 0 indicates soft sweep), f (frequency of selected mutation at sampling), g (number of generations since fixation), and s (selection coefficient).

### Caveats for Using Neutral Simulations to Normalize Real Data

In principle, one could use matched neutral simulations as a normalization baseline when analyzing real data. However, we can only recommend this approach when the populations being studied have very well characterized (1) joint demographic histories, (2) mutation rates, (3) and recombination rates, as a mismatch can skew the power and false positive rates of the statistic. To illustrate this point, we generated three mismatched sets of neutral simulations “Rand”, “Under”, and “Over” (see Methods) to use as a normalization baseline for our original simulations.

When neutral simulations are normalized with the correct demographic history, they approximately follow a standard normal distribution (Fig. S1 and Fig. S2), however the distribution of neutral scores gets badly distorted when one of the mismatched histories is used (Fig. S2). These distortions have practical consequences for making inferences. Power was calculated for XP-nSL using the window-based method described above but using each mismatched history as a normalization baseline (Fig. S3). The false positive rate for each scenario was also estimated by calculating the proportion of neutral simulations that are identified as under selection for each scenario (Table S2). Fig. S3A-B shows that for the “Rand” and “Under” normalization scenarios, power was greatly reduced across the whole parameter space. Only hard sweeps with the strongest selection coefficients post-fixation were likely to be identified. False positive rates for these scenarios were at 0 (Table S3), whereas for a matched history it was estimated at 9.908 × 10^-3^. Fig.S3C shows that, for the “Over” normalization scenario, power was uniformly excellent, however when analyzing neutral simulations, the false positive rate was estimated at 0.9538. This suggests that using such a badly matched demographic history for normalization creates a near complete inability to distinguish between neutral and selected regions.

### Identifying Evidence for Adaptation to High Altitude in Rhesus Macaques

Next, we analyzed the pair of rhesus macaque populations using XP-nSL, searching for evidence of local adaptation in the high-altitude population. Using the low-altitude population as the reference population, normalized *XPnS_L_*> 0 corresponds to longer and higher frequency haplotypes in the high-altitude population, with very large positive scores and clusters of large scores suggesting evidence for positive selection. Dividing the genome into 100 kb windows, the maximum XP-nSL score of that region is assigned to each gene in it (see Methods), thus multiple genes may have the same max-score by virtue of being in the same 100kb window. Genes overlapping multiple 100kb windows were assigned the top max-score among the windows.

Using the per-gene max-scores, PANTHER (Mi *et al*. 2019) gene ontology categories were tested for enrichment of high scores, where significance suggests an enrichment of signals of positive selection among genes involved. Significant terms related to regulation of ion transport and synaptic signaling (Table 2), each of which are affected by hypoxic conditions (Karle *et al*. 2004; Corcoran and O’connor 2013).

**Table 2.**
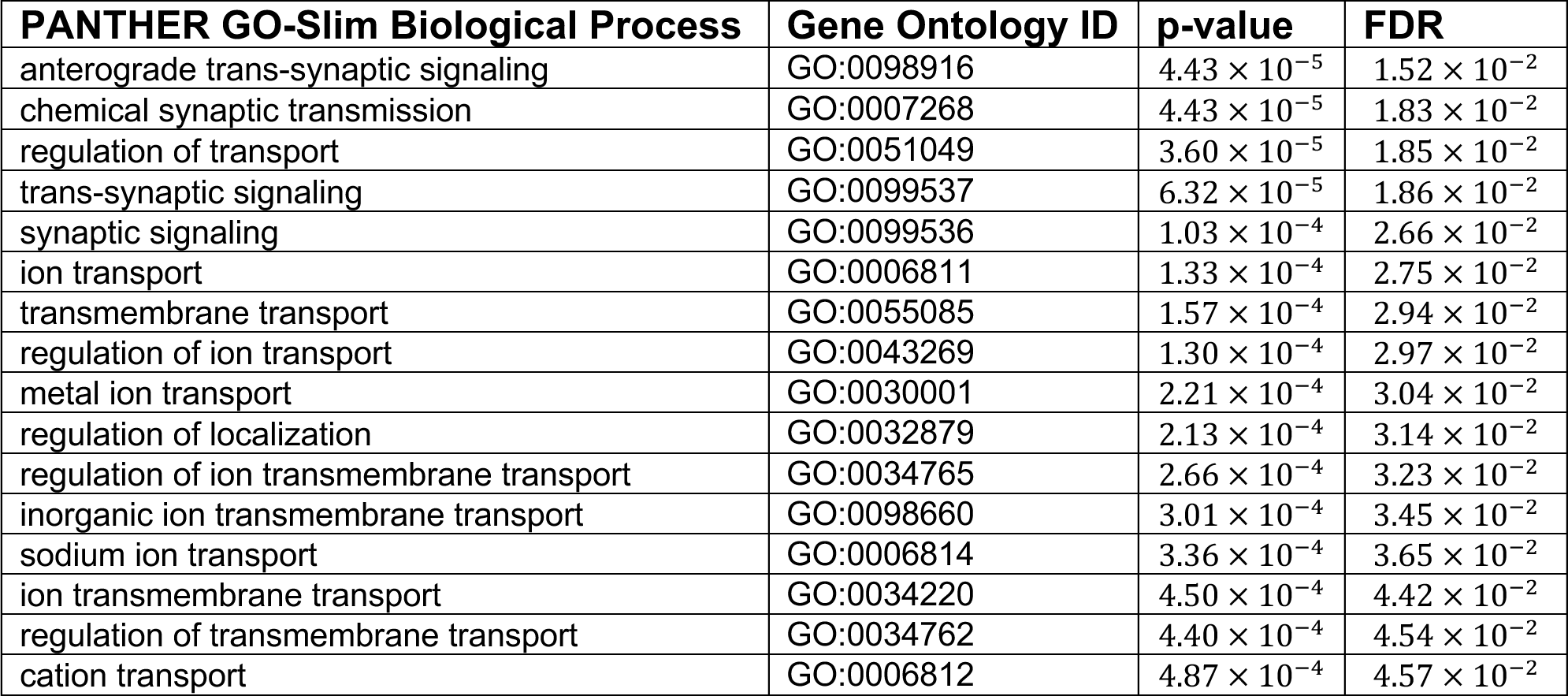
Gene ontology enrichment analysis results based on maximum XP-nSL scores per gene. Significant GO terms are enriched for high XP-nSL scores.

From the genomic regions that were identified to contain a high proportion of extreme positive scores (see Methods), 303 annotated genes were found across 270 regions. A permutation test (10,000 replicates) that randomly shuffles 270 100kb regions around the genome indicates that this is substantially fewer than one would expect by chance (*p* = 1.4 × 10^-3^; Fig. S5), indicating that the method is not simply randomly picking gene regions. These regions, their characteristics, and the genes contained therein are given in Table S2. A PANTHER (Mi *et al*. 2019) gene ontology overrepresentation test indicates a 9.04-fold enrichment of genes associated with monooxygenase activity (*FDR* = 4.47 × 10^-2^).

The monooxygenases in the selected regions include FMO2, FMO5, CYP2C8, CYP2C9, CYP2C93, and ENSMMUT00000011129. These genes are important for the metabolism of oxygen and the generation of reactive oxygen species (ROS) (Krueger and Williams 2005). Under the physiological stress of a low-oxygen environment, ROS levels increase and cause oxidative damage, and, in humans, long-term adaptation to high altitudes includes adaptation to oxidative damage (Janocha *et al*. 2017). Indeed, AOX1 is also identified in our top regions, mutations in which have been shown to affect ROS levels in humans (Foti *et al*. 2017).

The genome-wide top ten 100 kb windows based on the percentage of extreme XP-nSL scores are summarized in Table 3, and these windows overlap several genes, including EGLN1. The EGLN1 locus is directly adjacent to the single strongest selection signal identified in the entire genome (Fig. 6). This region has the third highest cluster of extreme scores (Table 3), contains the highest XP-nSL score in the entire genome (chr1:207,698,003, *XPnS_L_* = 6.54809), and contains six of the top ten genome wide XP-nSL scores (colored dark red in Fig. 6). EGLN1 is a regulator of oxygen homeostasis (To and Huang 2005) and is a classic target for adaptation to low-oxygen levels, having repeatedly been the target of adaptation in numerous organisms living at high altitude around the world (Bigham *et al*. 2009; Bigham *et al*. 2010; Jeong *et al*. 2014; Graham and Mccracken 2019). In addition to EGLN1, other genes related to lung function, oxygen use, and angiogenesis had evidence for local adaptation between low- and high-altitude populations: TRPM7, RBPJ, and ENSMMUT00000040566 (Table S2). TRPM7 downregulation in a hypoxia-induced rat model was associated with pulmonary hypertension (PAH) (Xing *et al*. 2019). ENSMMUT00000040566 is a MAPK6 ortholog, which interacts with EGLN3 (Rodriguez *et al*. 2016), and both it and RBPJ are involved in angiogenesis (Ramasamy *et al*. 2014).

**Figure 6.**
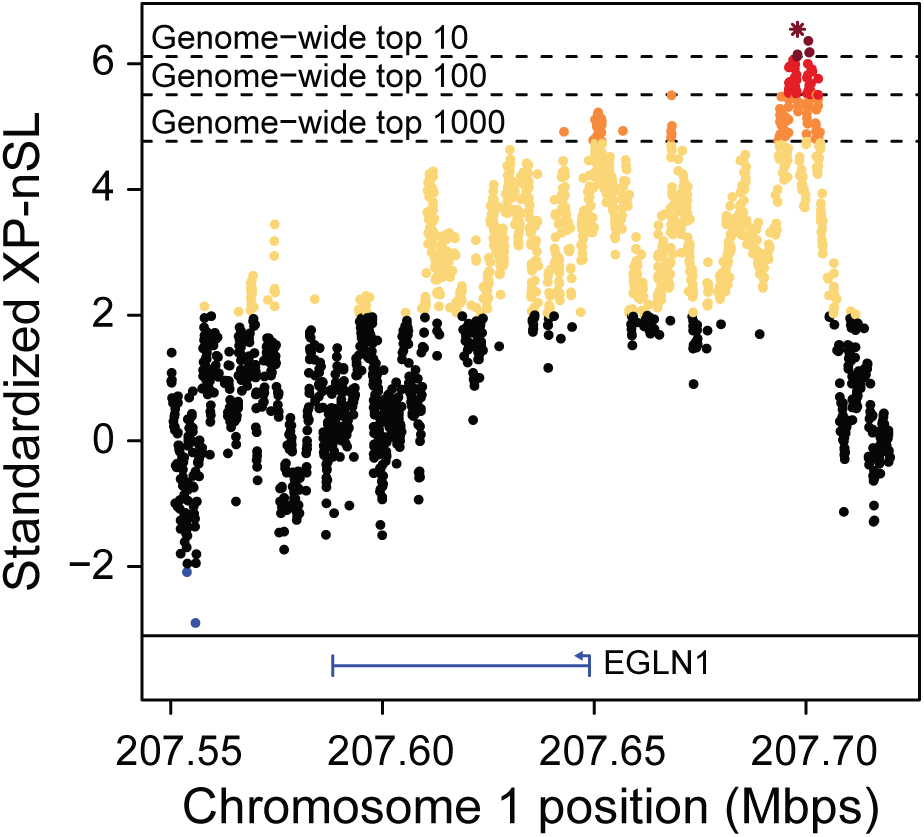
XP-nSL scores in the vicinity of the EGLN1 locus. This locus contains the genome wide top score (star) and six of the top ten genome wide scores (dark red).

**Table 3.** The top ten 100 kb genomic regions as ranked by percentage of scores greater than 2. Concatenated region represents the genomic region merged with adjacent top 1% regions. Max score represents the max XP-nSL score in the concatenated region. Genes gives all genes overlapping the concatenated region.

| Genomic Region | % > 2 | Flanking Regions | Concatenated Region | Max Score | Genes |
| --- | --- | --- | --- | --- | --- |
| *chr2:42300001-42400000 | 81.62% | 3 | chr2:42200001-42600000 | 5.42043 | STXBP5L |
| chr3:99100001-99200000 | 78.18% | 3 | chr3:99100001-99500000 | 5.54282 | - |
| **chr1:207600001-207700000** | **77.39%** | **1** | **chr1:207600001-207800000** | **6.54809<sup>†</sup>** | **EGLN1, TSNAX** |
| chr10:69700001-69800000 | 76.60% | 4 | chr10:69700001-70200000 | 4.89388 | PITPNB, ENSMMUT00000079195.1, TTC28 |
| chr13:64300001-64400000 | 74.01% | 1 | chr13:64300001-64500000 | 4.17728 | WDPCP, MDH1, ENSMMUT00000070474.1 |
| chr7:96400001-96500000 | 73.86% | 1 | chr7:96400001-96600000 | 4.58886 | FAM177A1, PPP2R3C, ENSMMUT00000039911.2, ENSMMUT00000027928.3 |
| chr10:70400001-70500000 | 70.24% | 0 | chr10:70400001-70500000 | 3.69192 | TTC28 |
| *chr2:42400001-42500000 | 65.86% | 3 | chr2:42200001-42600000 | 5.42043 | STXBP5L |
| chr7:120400001-120500000 | 62.54% | 0 | chr7:120400001-120500000 | 5.86114 | KIAA0586, ENSMMUT00000059161.1 |
| chr18:20600001-20700000 | 62.42% | 0 | chr18:20600001-20700000 | 5.37386 | RNF138 |

\*These regions are contained in the same concatenated region.
<sup>†</sup>Top genome-wide score. This region contains 6 of the top 10 genome-wide scores.

Due to the reduced oxygen levels at high altitudes, we expect genes involved in metabolism and respiration may be under positive selection. Indeed, MDH1 encodes a critical enzyme in the citrate cycle (Tanaka *et al*. 1996) and is found in the top ten genome-wide regions (Table 3). A paralog of MDH1, MDH1B, has been previously identified as a target of selection in humans living at high altitude (Yi *et al*. 2010). ACADM and COX15 are also found in putatively adaptive regions (Table S2). Mutations in and differing expression levels of ACADM are related to oxidative stress and mitochondrial dysfunction in human disease (Xu *et al*. 2018). COX15 is involved in oxidative phosphorylation (Alston *et al*. 2017), and cytochrome c oxidase (COX) genes have previously been identified as under selection in primates relative to other mammals (Osada and Akashi 2012).

High-altitude environments present a particular metabolic challenge to organisms that must maintain a stable internal body temperature (Rosenmann and Morrison 1974; Hayes and Chappell 1986; Chappell and Hammond 2004), a result of increased oxygen demand from aerobic thermogenesis conflicting with lower oxygen availability. It has been shown that highland deer mice have adapted by increased capacity to metabolize lipids compared to lowland deer mice (Cheviron *et al*. 2012), and previous studies in rhesus macaques have shown there may be drastic differences in diets between high- and low-altitude populations (Zhao 2018). Indeed, in the high-altitude macaque population studied here, genes related to lipid and fat metabolism (DOCK7, ST6GALNAC5, ANGPTL3, and ACACA) were found in putative adaptive regions. Across human populations, these genes are all responsible for varying blood levels of fatty acids (Guo *et al*. 2016; Dewey *et al*. 2017; Hebbar *et al*. 2018). Furthermore, ACACA has been shown to vary fatty acid blood concentrations and be differentially expressed in highland versus lowland swine populations (Shang *et al*. 2019).

STXBP5L also appears in the top ten regions (Table 3) and is involved in vesicular trafficking and neurotransmitter release (Kumar *et al*. 2015). As primate brains use large amounts of oxygen and energy to function (Osada and Akashi 2012) signatures of selection on neurological genes may be expected across populations living at altitudes with differing oxygen levels. In addition to STXBP5L, several genes related to neural development and synaptic formation (JAG2, TRPM7, DOCK7, NSG2, AUTS2) were identified (Table S2). JAG2 is involved in the Notch signaling pathway. While Notch signaling is involved in many developmental and homeostatic processes, its role in neuronal differentiation in the mammalian brain is notable in this context (Cardenas *et al*. 2018). TRPM7, in addition to its association with PAH, plays a role in hypoxic neuronal cell death (Aarts *et al*. 2003).

DNA damage, including double strand breaks and pyrimidine dimerization, can manifest as a result of oxidative stress (Ye *et al*. 2016) or increased exposure to UV radiation (Zhang *et al*. 2000; Greinert *et al*. 2012), both of which increase at high altitudes. Three DNA damage repair genes are in our set of genes identified in positively selected regions. The ring finger protein RNF138 is in our list of top ten genomic windows (Table 3) and has been shown to promote DNA double-strand break repair (Ismail *et al*. 2015). Furthermore, the DNA polymerase POLH also appears in a putatively adaptive region (Table S2) and is known to be able to efficiently bypass pyrimidine dimer lesions (Zhang *et al*. 2000). Finally, PAXIP1 appears in Table S2 as well and has been shown to promote repair of double strand breaks through homologous recombination (Wang *et al*. 2010).

Interestingly, within the putatively selected region that includes PAXIP1 is an uncharacterized long non-coding RNA (lncRNA), ENSMMUT00000081951, that was recently annotated in the rheMac10 genome build. This lncRNA has high sequence similarity to a lncRNA on the same synteny block in humans called PAXIP1-AS1. In human pulmonary artery smooth muscle cells, knockdown of PAXIP1-AS1 leads to an abnormal response to PAH where migration and proliferation of cells is reduced, and overexpression of PAXIP-AS1 leads to apoptosis resistance (Jandl *et al*. 2019). Although, we note that any link between PAH and ENSMMUT00000081951 in rhesus macaques is highly speculative at this point.

## Discussion

When populations split and migrate, they may adapt in different ways in response to their local environments. Genetic adaptations that arise and sweep through the population leave a characteristic genomic pattern of long haplotypes of low diversity and high frequency. We develop a two-population haplotype-based statistic, XP-nSL, to capture these patterns. With good power to detect hard and soft sweeps that occur in one population but not another, XP-nSL can identify positively selected regions of the genome likely the result of local adaptation. We apply this statistic to genomes sampled from a pair of wild-born populations of Chinese rhesus macaques, inferred to have diverged approximately 9500 generations ago, one of which lives at high altitude in far western Sichuan province, and the other that lives close to sea level.

Life at high altitude presents extreme biological challenges, including low atmospheric oxygen, increased UV exposure, harsh winters, and reduced nutrition availability, which create strong selection pressure. Organisms that survive and persist are likely to be carrying genetic mutations that confer an advantage for living in such harsh environments. Common targets for adaptation to such an environment include genes related to hypoxia, regulation of ROS, DNA damage repair, and metabolism (Cheviron and Brumfield 2012; Witt and Huerta-Sanchez 2019; Storz and Cheviron 2021). Indeed, in the high-altitude macaque population, we identify a strong signal of positive selection at the EGLN1 locus (Fig. 6), a classic target for adaptation to low-oxygen environments, in addition to 302 other genes, many of which are related to the myriad environmental selection pressures expected in high-altitude environments. As has been suggested previously for other organisms (Cheviron and Brumfield 2012; Bigham and Lee 2014; Simonson 2015; Witt and Huerta-Sanchez 2019; Storz and Cheviron 2021), these results suggest that, rather than a single adaptive mutation at a single locus, adaptation to this extreme environment by rhesus macaques is polygenic and acts through multiple biological systems.

## Acknowledgments

The authors would like to thank members of the Stevison Lab for helpful discussions, Lawrence Uricchio for helpful comments on early versions of the manuscript, and two very helpful anonymous reviewers. This work was supported by start-up funds from the Department of Biological Sciences at Auburn University (LSS) and the Department of Biology at the Pennsylvania State University (ZAS). ZAS was partially supported by NSF-DEB EAGER No. 1939090 (LSS). Portions of this research were performed on the Pennsylvania State University’s Institute for Computational Data Sciences’ Roar supercomputer.

## Author Contributions

ZAS and LSS conceived of the study. ZAS performed all simulations and genomic analyses and implemented novel statistics. ZAS, TEN, and NPB characterized gene functions and ontologies with contributions from LSS. ZAS wrote the manuscript with contributions from NPB, TEN, and LSS. All authors read and approved of the manuscript.

## Data Accessibility

Macaque whole genome VCFs are available at http://dx.doi.org/10.5524/100484. Selection scan data available at https://doi.org/10.5061/dryad.kkwh70s40.

---

**Table S1.** Population classification by individual ID.

*See TableS1.xlsx*

**Table S2.** List of genomic windows with the top 1% highest fraction of extreme XP-nSL scores.

*See TableS2.xlsx*

**Table S3.** Estimates of false positive rates for various demographic history normalization scenarios.

| Demographic History | Estimated False Positive Rate |
| --- | --- |
| Matched | $9.908 \times 10^{-3}$ |
| "Rand" | 0 |
| "Under" | 0 |
| "Over" | 0.9538 |

**Table S4.** Bin boundaries (# of sites), number of windows per bin, and top 1% thresholds for the XP-nSL analysis of rhesus macaques.

| Bin Boundaries (min # sites, max # sites) | # Windows in Bin | Top 1% Threshold (Fraction of Scores > 2) |
| --- | --- | --- |
| (11, 731) | 2672 | 0.346755 |
| (732, 987) | 2680 | 0.262945 |
| (988, 1155) | 2676 | 0.207089 |
| (1156, 1285) | 2675 | 0.196678 |
| (1286, 1390) | 2663 | 0.226605 |
| (1391, 1484) | 2687 | 0.19274 |
| (1485, 1576) | 2665 | 0.163597 |
| (1577, 1681) | 2648 | 0.169275 |
| (1682, 1824) | 2666 | 0.164025 |
| (1825, 5766) | 2666 | 0.153176 |

**Figure S1.**
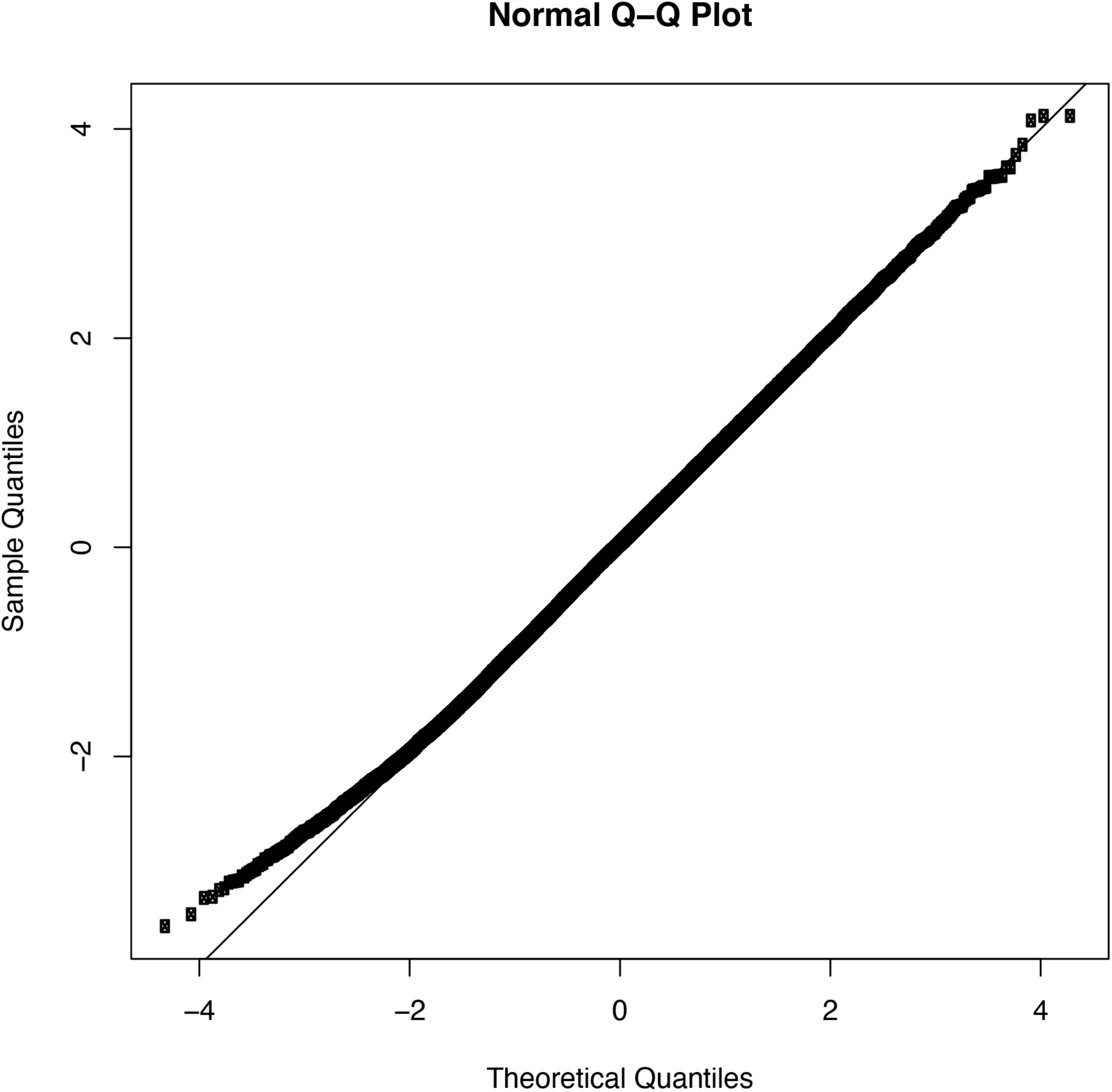
A normal quantile-quantile plot of neutral XP-nSL scores showing generally good adherence to a standard normal distribution. Due to autocorrelation along the genome, only every 1000^th^ score is plotted.

**Figure S2.**
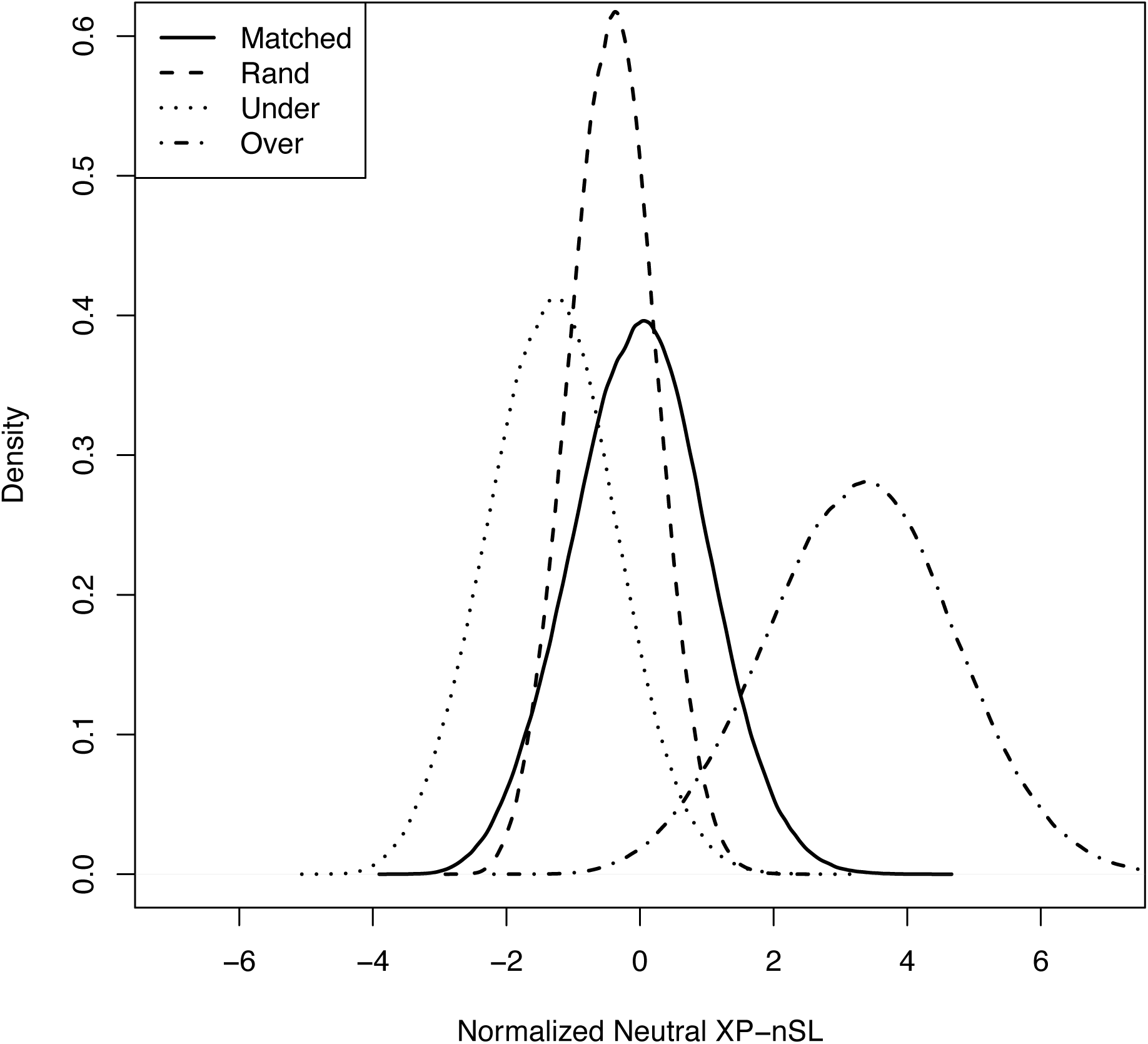
The distribution of neutral XP-nSL scores normalized with a matched demographic history (solid line), normalized with the “Rand” demographic history (dashed line), normalized with the “Under” demographic history (dotted line), and normalized with the “Over” demographic history (dash-dot line). Normalizing with the wrong demographic history can dramatically shift the distribution of neutral XP-nSL scores.

**Figure S3.**
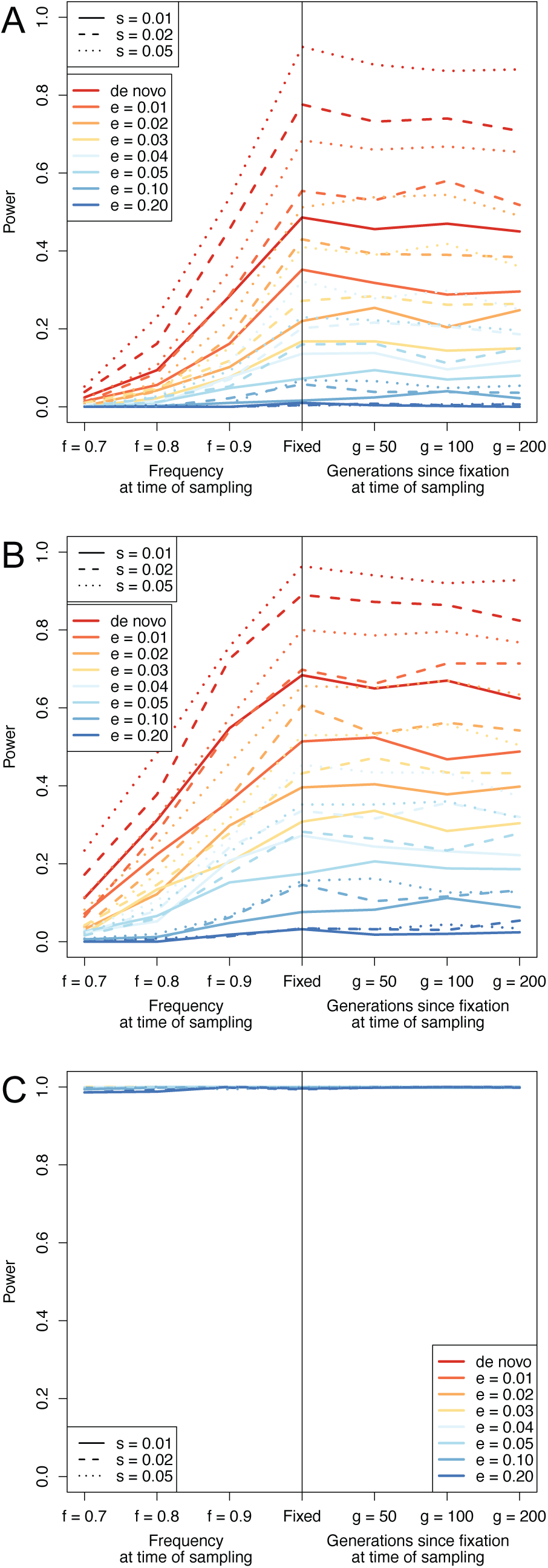
XP-nSL power using mismatched demographic histories for normalization. (A) Using the “Rand” history. (B) Using the “Under” history. (C) Using the “Over” history.

**Figure S4.**
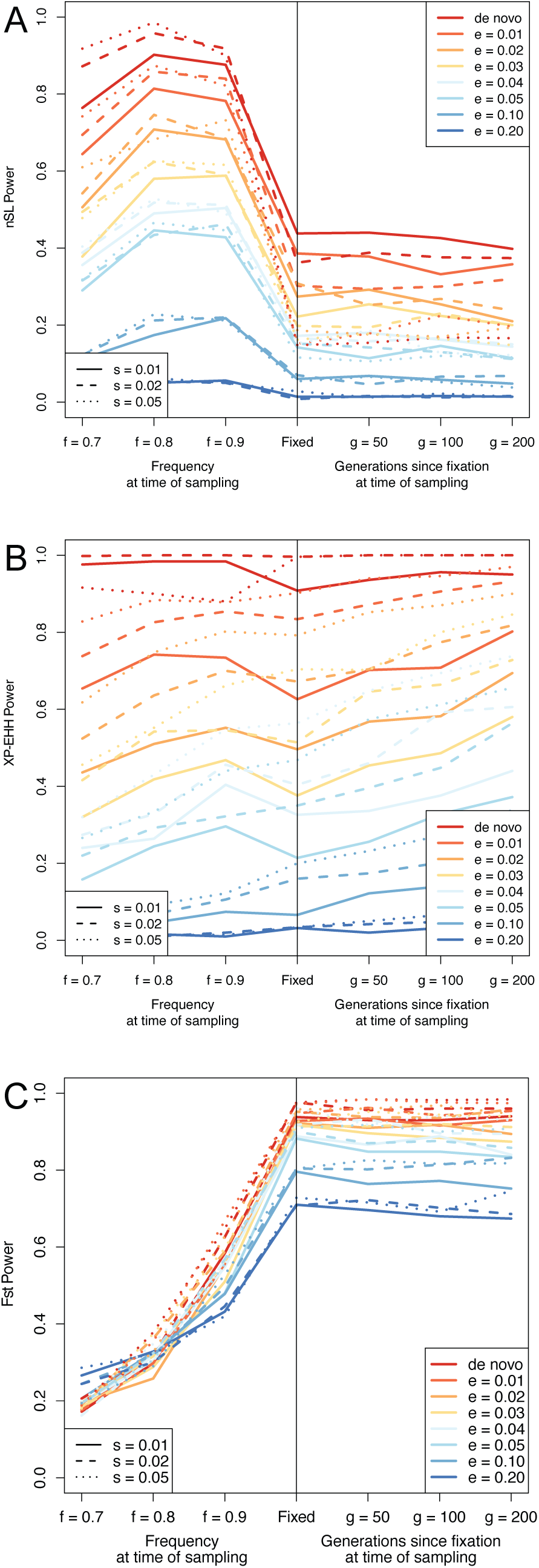
Power curves for (A) nSL, (B) XP-EHH, and (C) F_ST_. The parameters are e (frequency at which selection begins, e > 0 indicates soft sweep), f (frequency of selected mutation at sampling), g (number of generations since fixation), and s (selection coefficient).

**Figure S5.**
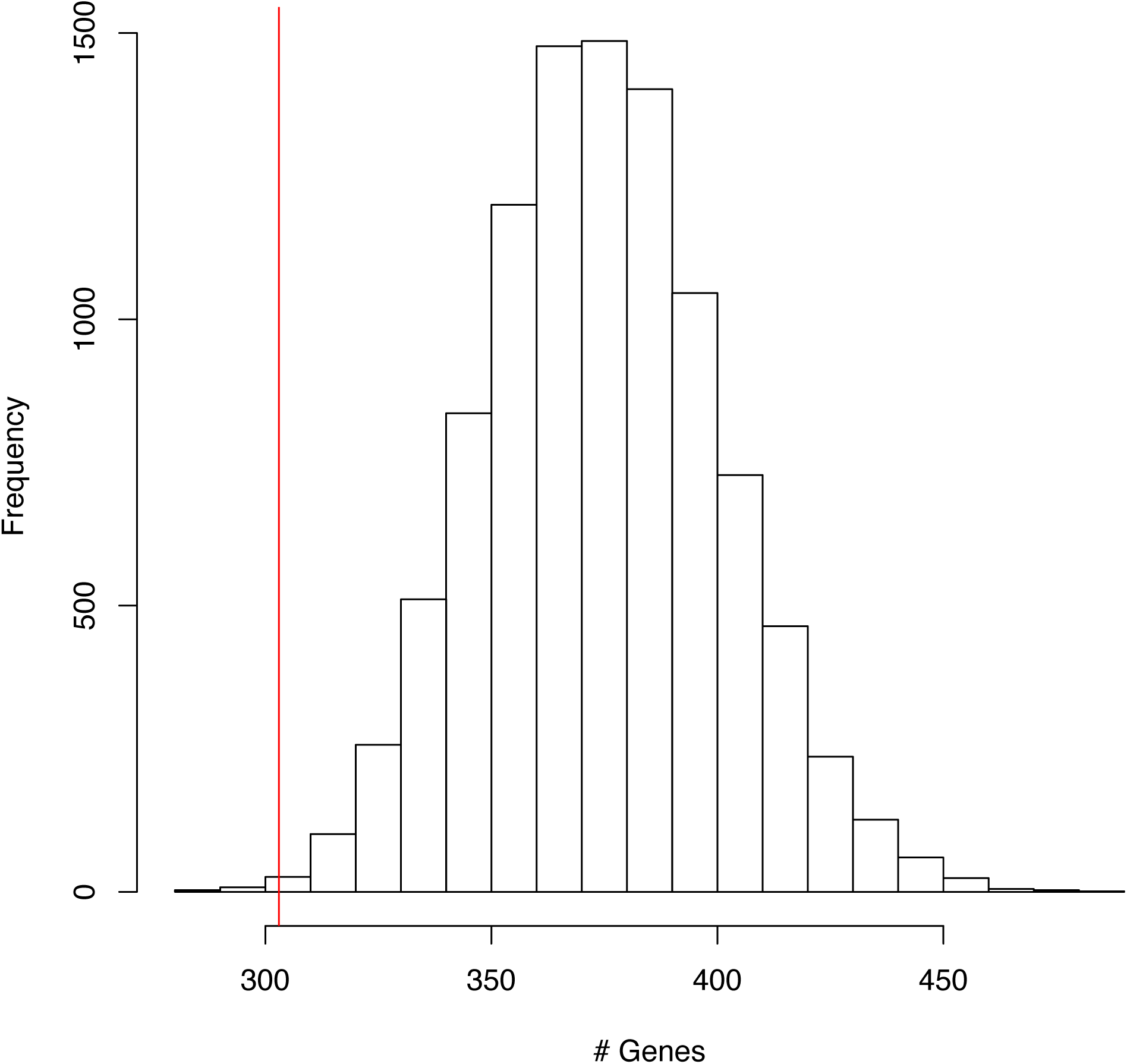
A permutation test (10,000 replicates) that shuffles 270 100kb regions around the macaque genome and counts the number of unique genes overlapping. The red vertical line marks the 303 genes found in the real data analysis. The probability of observing 303 or fewer genes is 1.4 × 10^-3^, indicating the analysis is not randomly choosing gene regions.

**Figure S6.**
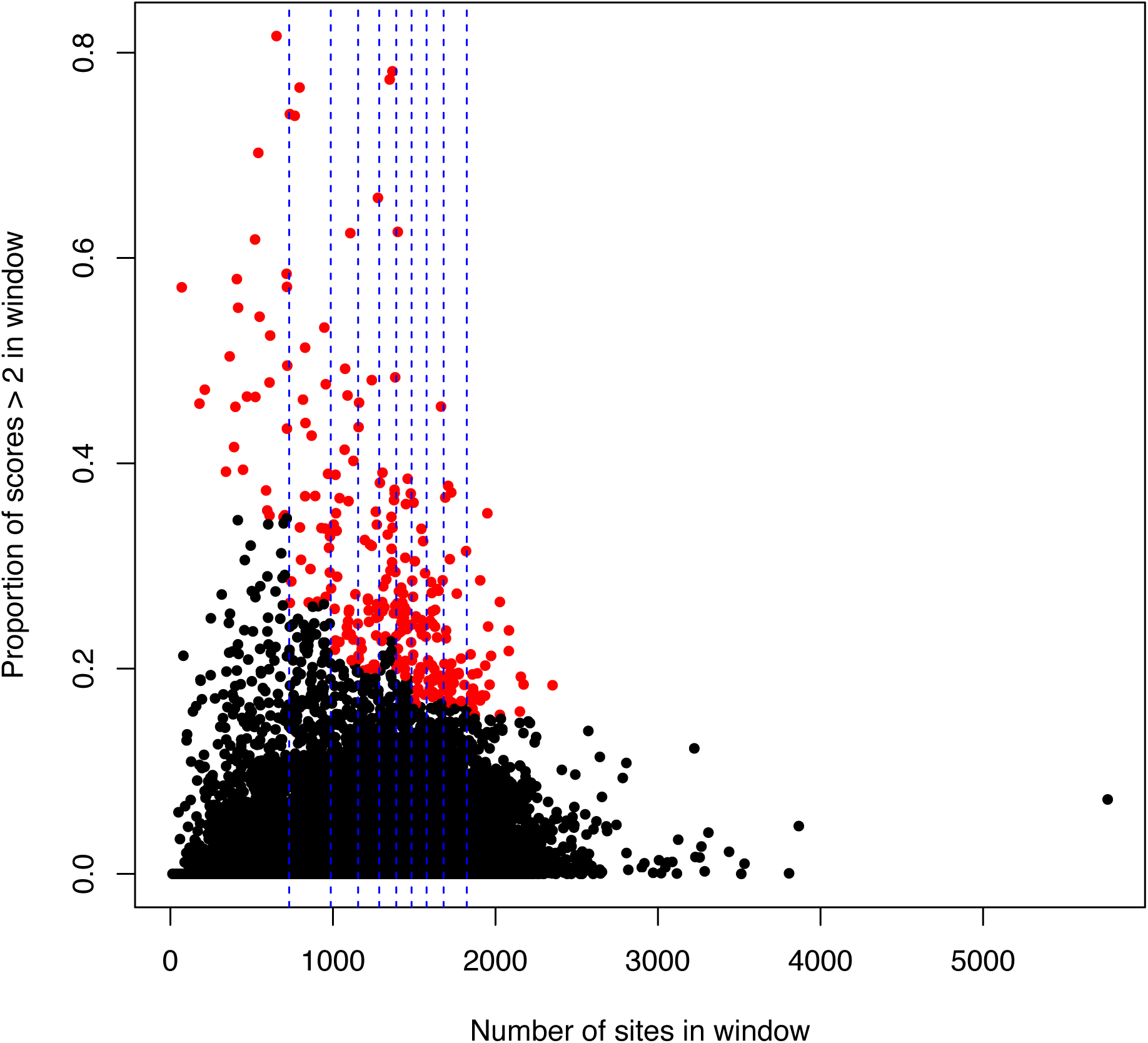
Proportion of scores > 2 versus number of sites in window. Blue vertical dashed lines indicate bin boundaries. Each circle is a window, red dots indicate a proportion of scores > 2 beyond the 1% threshold for that bin.

**Figure S7.**
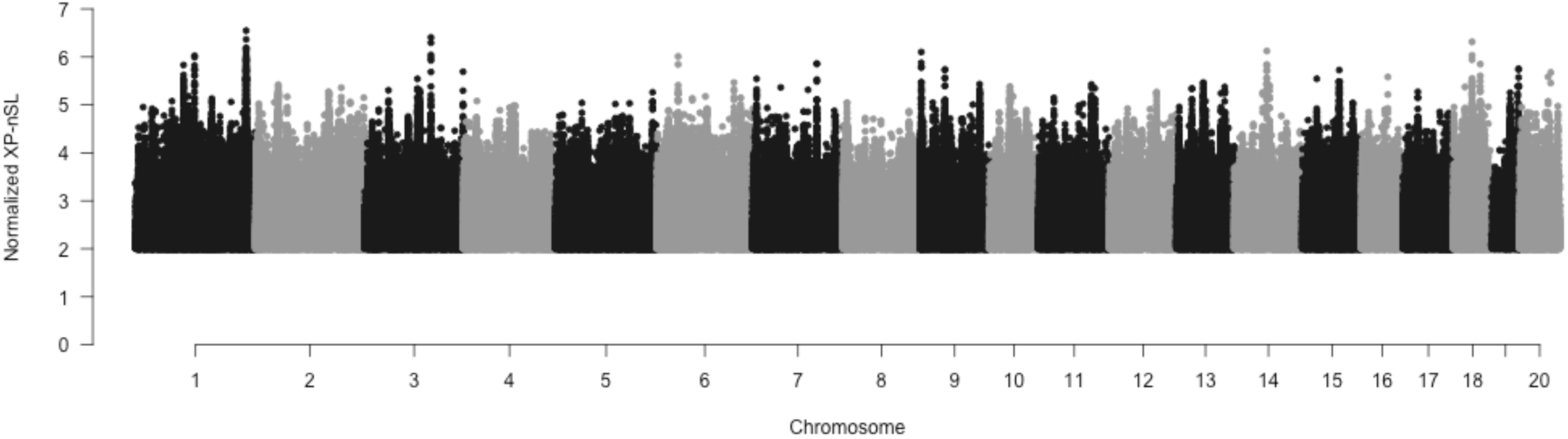
A Manhattan plot of normalized XP-nSL scores across the genome. Due to a very large number of points, only scores > 2 were plotted.

